# EnviroAmpDesigner – a tool for designing high specificity multiplex PCR primer panels for detecting and subtyping a target organism in environmental surveillance samples

**DOI:** 10.1101/2025.04.10.648097

**Authors:** Anton Spadar, Jaspreet Mahindroo, Catherine Troman, Michael Owusu, Yaw Adu-Sarkodie, Ellis Owusu-Dabo, Dilip Abraham, Blossom Benny, Karthikeyan Govindan, Venkata Raghava Mohan, Zoe A. Dyson, Nicholas Grassly, Kathryn E. Holt

## Abstract

**Background:** Amplicon sequencing is a popular method for understanding the diversity of bacterial communities in environmental or similar samples as exemplified by 16S rRNA sequencing. This approach has been extended into multiplex amplicon sequencing in which multiple targets are amplified in the same PCR reaction such as virus sequencing using tiled amplicons. Multiple tools exist to design PCR primers, and some support design of multiplex panels. However, despite increasing interest in the use of environmental or wastewater sampling for detecting and typing specific antimicrobial resistant (AMR) bacteria such as typhoid or cholera agents, we were unable to find a tool for designing a multiplex PCR panel for environmental samples that not only focused on detection of a specific organism, but also on amplifying lineage-specific or AMR-associated alleles of the target organism while minimising amplification of off-target genomes present in the sample. We found that existing tools either depend on the target organism being very distinct from the rest of the organisms in the sample, or focus on detection rather than genotyping of the organism of interest. We have developed EnviroAmpDesigner (v0.1.3, DOI: 10.5281/zenodo.14967337) to fill this gap, which we used to design a multiplex amplicon panel for detection, genotyping and AMR profiling of the typhoidal pathogens *Salmonella enterica* serovars Typhi and Paratyphi A. EnviroAmpDesigner can design amplicons for both short and long-read sequencing. The software first identifies single nucleotide polymorsphisms (SNPs) that distinguish individual genotypes of the target pathogen (in our case, *S*. Typhi). It then identifies SNPs that distinguish the target organism from others. Finally, the software designs primers that simultaneously target genotyping SNPs and have at least one primer with a 3’ end at each SNP that distinguishes the target organism. The purpose of the latter condition is to increase the specificity of the primers, to minimise amplification of homologous sequences in non-target organisms. While our use case focuses on *S*. Typhi and Paratyphi A, the tool is organism agnostic and should work for any haploid organism. The tool is freely available via Bioconda and GitHub (https://github.com/AntonS-bio/EnviroAmpDesigner).

**Impact statement:** We have developed EnviroAmpDesigner (v 0.1.3) software to design a multiplexed panel of long-read amplicons for detection and lineage typing of bacteria in environmental samples. We apply this tool to *Salmonella* Typhi and Paratyphi A, the bacterial agents of typhoid fever, but the tool is organism agnostic. The tool is computationally efficient and can be used on mid-range laptop. It has potential to support more widespread deployment of sequencing-based surveillance for bacterial pathogens in environmental and wastewater samples.

**Data Summary:** For the purposes of designing primers for *S*. Typhi and Paratyphi A we used 13,134 samples from the Global Typhoid Genomics Consortium data (target organisms) (1) and 1,810 publicly available Enterobacterales genomes from NCBI’s RefSeq database (representing off-target related organisms) (2). To validate the primers, we applied an amplicon sequencing protocol (3) to DNA extracted from two *S*. Typhi isolates (NCBI accessions SRR5949979 and SRR7165748) provided by Satheesh Nair (UKHSA) (4), and a pooled DNA sample of American Type Culture Collection (ATCC) isolates *S*. Paratyphi A (ATCC 9150D), *S*. Paratyphi B (ATCC-BAA-1250D), *S*. Paratyphi C (ATCC-BAA-1715D), *Aeromonas hydrophila* (ATCC-7965D), *Klebsiella pneumoniae* (ATCC-BAA-1706D), and *Citrobacter freundii* (ATCC-8090D) chosen for their close relationship to *S*. Typhi. The resulting amplicon sequence data was deposited in European Nucleotide Archive (project accession: PRJEB81565).

## Introduction

Detection and typing of pathogens direct from wastewater or environmental samples has gained popularity as a public health surveillance measure, allowing both the monitoring of pathogenic organisms circulating amongst human populations (shed into wastewater) as well as routes of pathogen dissemination and exposure (e.g. presence in surface water or drinking water) (5). A common strategy is to employ PCR to amplify sequences unique to a target organism and use qPCR to confirm and quantify amplification (6, 7). However, this can lead to false positives especially in the context of samples from complex sources such as environmental surveillance, where the target pathogen sequence may be in very low abundance and highly similar to other non-target sequences in the same sample. One solution is to combine PCR with sequencing, whereby amplicons are sequenced to confirm their identity and distinguish on-target from off-target amplification. Amplicon sequencing also offers the opportunity to ‘subtype’ target organisms based on sequence variation within the amplicons. This approach has been used for poliovirus surveillance (6). In the microbiome field, 16S rRNA amplicon sequencing is a well-established approach to taxonomic profiling of bacteria present in mixed samples, whereby a single set of primers amplifies rRNA from diverse species and the amplicon sequences are classified taxonomically to provide insight into the composition of the sample. Multiplexing is also possible (5), whereby multiple targets are amplified in the same PCR reaction (via inclusion of multiple primer pairs), and the resulting amplicon sequenced to identify which targets were present, any genetic variation within them, and their relative abundance (8, 9). Inclusion of primers targeting regions relevant to antimicrobial resistance (AMR) or pathogenicity mechanisms can offer additional insights into resistance or virulence of the target pathogen. Long-read sequencing platforms such as Oxford Nanopore Technologies (ONT) allows the multiplexing of long amplicons (several kbp long) with substantial discriminatory power, using portable devices with low capital expenditure (10, 11).

Multiplex amplicon sequencing via ONT therefore offers a very flexible and generalisable approach for detection and typing of pathogenic organisms in mixed samples, that is in principle feasible for wastewater and environmental surveillance in a broad range of settings including low-resource laboratories or field settings. However, designing primer sets for such applications can be challenging, due to the requirements for both high sensitivity (problematic for environmental or similarly complex samples) and high specificity (to avoid the PCR reaction being overwhelmed by off-target sequences). For example surveillance of typhoid fever, and particularly AMR infections, is challenging in many countries where typhoid fever is endemic due to a lack of blood culture to isolate the pathogen (*Salmonella enterica* serovar Typhi, *S*. Typhi) and test its antimicrobial susceptibility. We were therefore motivated to design a multiplex amplicon sequencing assay to detect *S*. Typhi, and also discern common AMR-associated *S*. Typhi lineages and AMR determinants. This is particularly challenging as *S*. Typhi is expected to be present in a dormant state and at very low abundance(12, 13), and genetically close relatives are expected to be present in much greater abundance (including other more common *Salmonella enterica* serovars, and other members of the bacterial family *Enterobacteriaceae* which include many environmental bacteria)(14).

To our knowledge, there are no actively supported tools for designing primers for genotyping a single organism in the sample. We have found that some online tools are no longer available (15, 16), while the RUCS software (17) was functioning, but was no longer maintained, had a very long runtimes and was not designed for multiplex PCR. We therefore created EnviroAmpDesigner to fill this gap It uses user-provided VCF files and genomic sequences of target and non-target organisms to design a set of multiplex PCR primers to target single nucleotide polymorphisms (SNPs) that distinguish user-defined genotypes of the target organism while avoiding amplifying non-target organisms.

## Theory and implementation

### Overview

EnviroAmpDesigner creates a list of primer pairs for multiplex PCR that, when the resulting amplicons are sequenced, can detect and type a target organism in a DNA sample containing large number of different organisms. We define target organism as the organism whose presence the user wants to detect in the sample, and non-target organisms as all other organisms. We define genotypes as a set of user supplied genotype labels (i.e. lineage designations) for the target organism. We define target genotypes as a subset of genotypes whose presence the user wants to specifically detect using multiplex PCR. The non-target genotypes are all genotypes of the target organism that do not need to be specifically differentiated by the assay **(Fig. 1)**.

**Figure 1.**
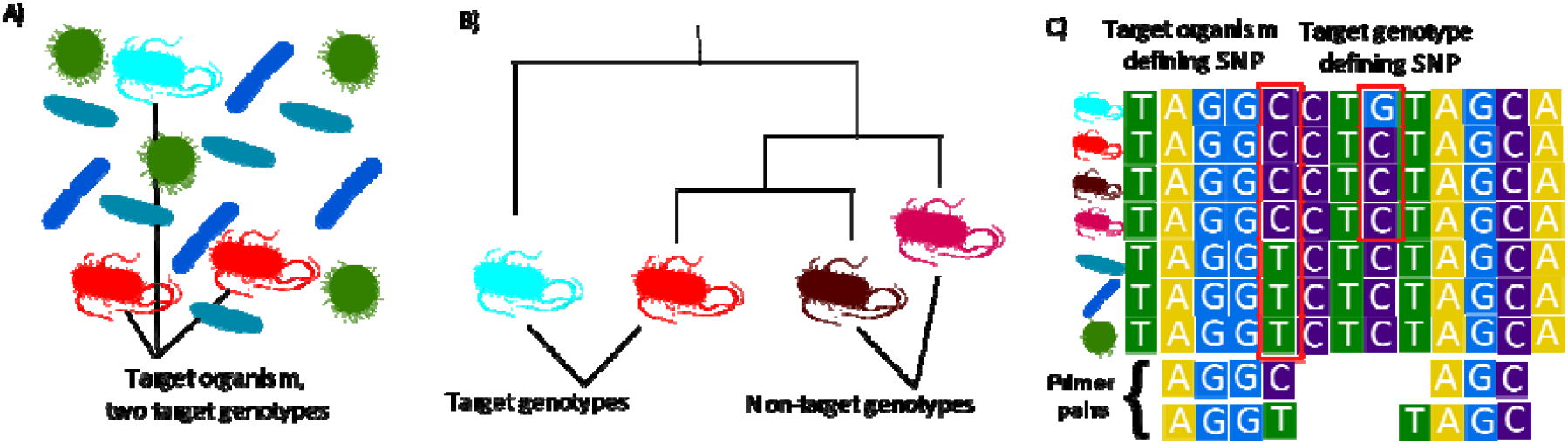
Schematic illustrating the purpose and strategy of EnviroAmpDesigner. The purpose of EnviroAmpDesigner is to design primers for multiplex PCR and sequencing to detect and type a target organism in a mixed sample. It aims to generate primers whose sequences can be used to confirm the presence of the target organism and distinguish target genotypes (i.e. lineages) of that organism, while avoiding amplification of non-target organisms. (A) Organisms present in a mixed sample; this example includes 2 different genotypes of the target organism (red, cyan) and several non-target organisms. (B) Phylogenetic tree showing all 4 genotypes defined for the target organism, highlighting 2 target genotypes whose presence the primers should aim to specifically identify. (C) Sequence alignment highlighting the ideal scenario for primers that EnviroAmpDesigner aims to identify; specifically, a target genotype-defining SNP within X bp of an organism-defining SNP. Primers are designed such that at least one has an organism-defining SNP as its final 3’ nucleotide (to reduce binding to non-target organisms), and the target-defining SNP lies within the resulting amplicon (allowing the allele to be determined by sequencing the amplicon).

EnviroAmpDesigner works in three stages **(Fig. 2)**. The first stage identifies target organism SNPs that separate target from non-target genotypes **(Fig. 1)**. To do this, EnviroAmpDesigner needs to have a user provided set of VCF files which contain variant calls for target organism SNPs against a reference genome, for a large set of isolates representative of population variation in the target organism. To create these VCF files, the user should select a reference genome of the target organism, and align a large set of target organism whole genome sequencing (WGS) read sets to it (e.g. using XX) and call variants (e.g. using bcftools). Ideally each target genotype (lineage) label should be represented by WGS libraries from multiple biological replicates of that genotype. The specific choice of reference genome, for the purpose of amplicon design, is not important but needs to be consistent across all VCFs. EnviroAmpDesigner requires genotype labels for all samples in the VCF files, supplied as a comma or tab delimited metadata file. The only requirement for this metadata file is that the first column contains the names of the corresponding VCF file (without .vcf) and that some column contains the genotype label. In addition, users must supply a file indicating which genotypes should be targeted and specifying the hierarchical relationships between genotypes (see **Fig. 3**).

**Figure 2.**
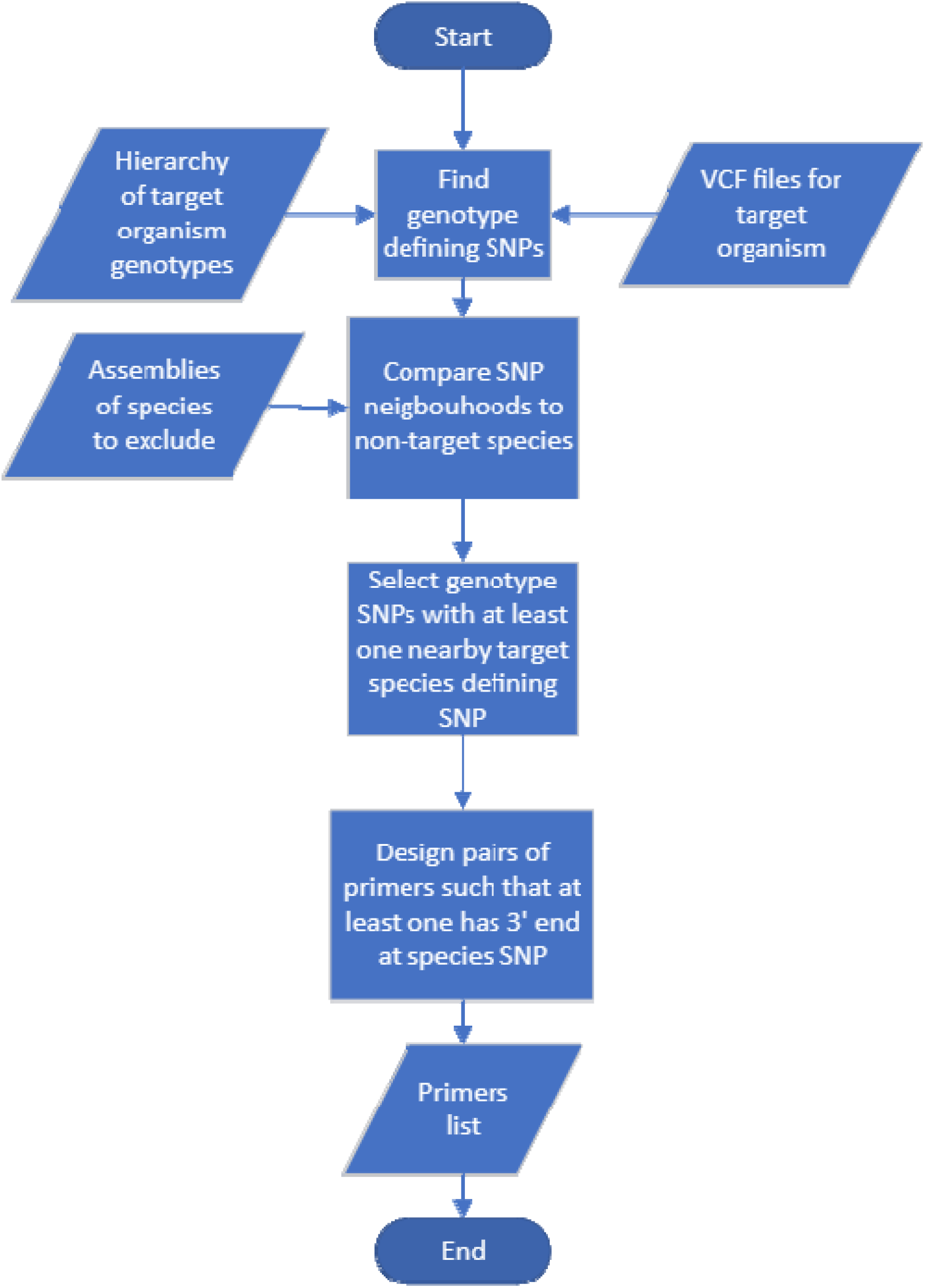
Simplified program workflow. Detailed version is in Supplementary Figure A and B.

**Figure 3.**
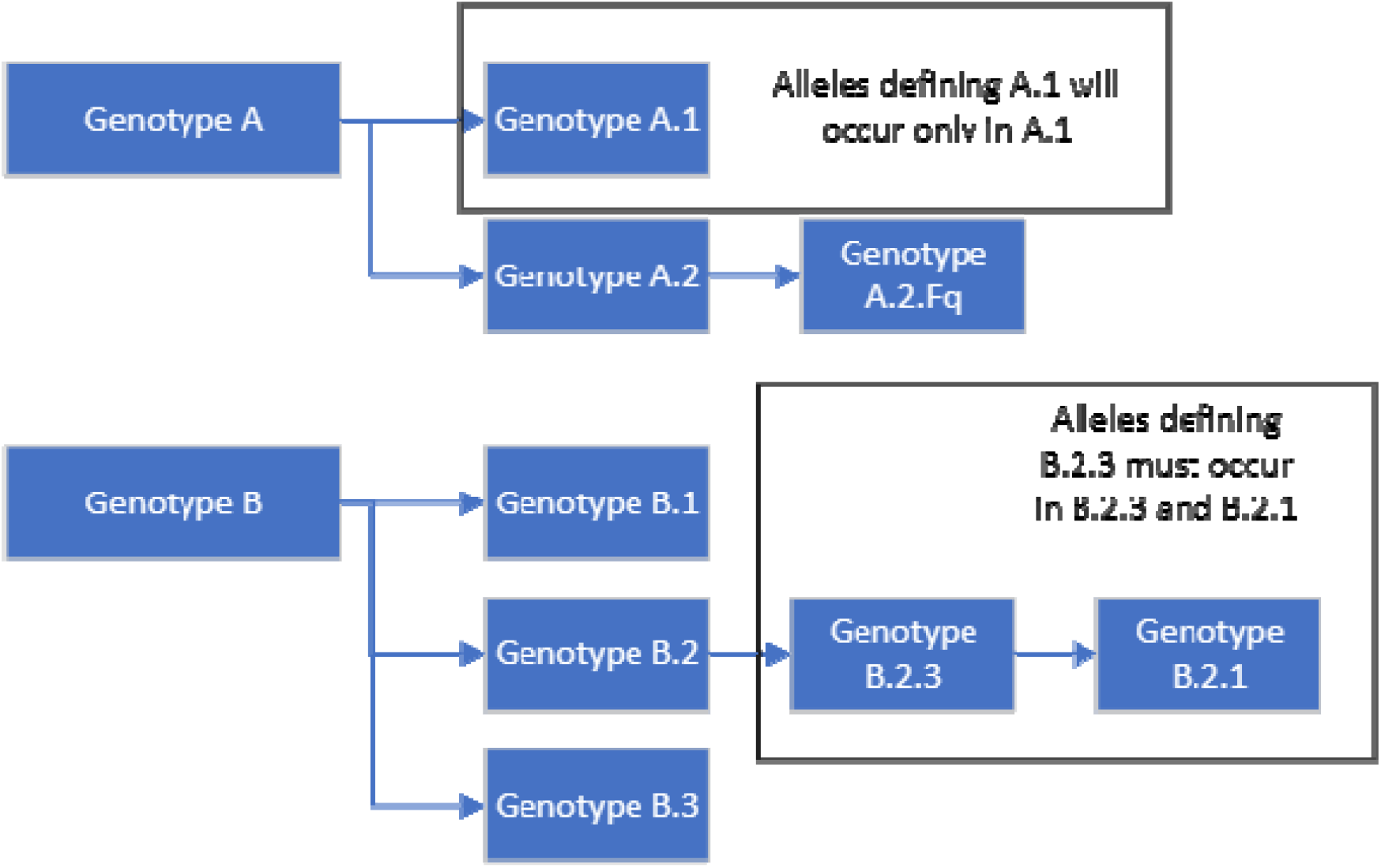
Handling of hierarchical genotypes or lineage labels. Lineage genotyping schemes are typically hierarchical, with genotypes subdivided into higher-resolution genotypes as pathogens evolve and diversify into subgroups. If a target genotype label has subtypes, primers need to be designed such that all subtypes can be detected. This hierarchical structure therefore needs to be given as input to EnviroAmpSeq, so that it can parse the relationships between genotype labels provided for each input VCF (while some schemes aim to indicate relationships between genotypes in the names, there are sometimes exceptions, and there is no consistent nomenclature used across schemes, so this must be directly specified). (A) Schematic showing hierarchical relationships for an example genotype scheme. (B) Representation of the genotype hierarchy in the format required by EnviroAmpSeq. In the example given here, genotype label B.2.1 is inconsistent with the rest of the scheme because it is a sub-genotype label of B.2.3 while A.2.Fq does not follow numeric format of the rest of the scheme. The genotypes relationship is a required input because EnviroAmpDesigner needs to know that B.2.1 is a sub-genotype label of B.2.3.

In the second stage, EnviroAmpDesigner identifies nucleotides that differentiate target from non-target organisms **(Fig. 1, 2)**. To do this, the tool requires whole genome assemblies of non-target organisms. For each SNP selected at stage one EnviroAmpDesigner compares the SNP neighbourhood on the target organism reference genome to all non-target organism genomes (see details below). Based on this comparison, the tool searches for target genotype-defining SNPs with at least one proximate SNP, that differentiates target and non-target organisms. Maximum proximity is defined as distance on the reference genome defined by user in configuration file. **(Fig. 1)**.

In the third stage, EnviroAmpDesigner tests all possible primer pairs in the neighbourhood of each target genotype-defining SNP, searching for those where at least one primer in each pair has as the last based at the 3’ end a SNP that distinguishes target and non-target species **(Fig. 1)**. This is a key strategy to minimise primer binding to closely related non-target organisms, aiming to reduce the PCR reaction being overwhelmed by amplification of non-target DNA.

Typical EnviroAmpDesigner analysis requires a large number of inputs and significant run time. We have introduced several input validation steps to ensure that input related errors are flagged before the start of analysis and accompanied by useful error messages. The only non-standard input file is the genotypes hierarchy which tells it which genotypes should be targeted, and also what are the sub-genotypes of these genotypes **(Fig. 3)**. For flexibility, we provide two options to specify this. If user already has structure of genotypes in JSON file **(Supp. File 1)**, this can be used as input along with plain text file which lists the target genotypes. Alternatively, if user does not have hierarchy in JSON file, they can provide a single tab delimited file in which every line represents a target genotype. In this case, each line should start with an id of the target genotype label and the rest of the line should be a tab-delimited list of all the subgenotypes of this genotype. This means that some lines are going to be very long, and others might have a single value in them. For example, **Supp. File 2**, means that user wants to target genotypes A, A.1, B, and B.2.3 from **Fig. 3**.

### Stage 1: SNP identification

In the first stage EnviroAmpDesigner designs multiplex PCR primers that target SNPs that split samples into subsets based on labels provided in a column of the metadata file **(Fig. 2)**. The primary use case is to target genotype-defining alleles (in which case the genotypes hierarchy file must also be provided), but in principle any other column from the metadata file can be targeted instead (e.g. an antimicrobial resistance or virulence phenotype).

EnviroAmpDesigner loads VCF files into memory, and stores them as a set of SNP objects and Sample objects **(Supp. Fig. 1)**. This approach means that multiple occurrences of the same SNP across samples requires a single object in memory and each sample merely points to SNP objects. This reduces memory footprint by an order of magnitude compared to representing the same data as matrices or data frames. Importantly, EnviroAmpDesigner disregards positions with multiple alternative alleles because they have strong likelihood of being under selective pressure and thus unstable relative to the target subgroup.For each SNP in the VCF files EnviroAmpDesigner determines if there is a target genotype label that is defined by this SNP **(Fig. 1, Supp. Fig. 1)** using the full set of VCF input and information on relationship between genotypes **(Supp. File 1, 2)**. Such genotype label defining SNPs are kept for further analysis, and the rest are discarded. The target is considered defined by a SNP if the SNP has specificity and sensitivity at least as high as set in config file **(Supp. File 3)**, the suggested default being 98%.

In some use cases the number of target genotypes can be very high. For example, in our use case of *S*. Typhi we targeted 17 genotype labels. To minimise the number of primer pairs and thus amplicons needed to cover the targets, by default EnviroAmpDesigner prioritises genotype-defining SNPs that lie close enough to another genotype-defining SNP to be captured by the same amplicon. The acceptable proximity is defined as a pair of SNPs within a given nucleotide distance of each other (parameter “flank_len_to_check” in the config file, suggested default being 1000 which is suitable for current ONT technology). However, there are two cases in which all genotype-defining SNPs may be taken to the next stage: the genotype label has either <10 defining SNPs, or it is listed in field “gts_with_few_snps” of the config file. Using this approach, for our use case we were able to halve the number of amplicons needed to differentiate all target genotypes; though the benefit of this approach will vary depending on the target organism and genotypes.

### Stage 2: Homologues identification

In the second stage, EnviroAmpDesigner identifies the target organism-defining SNPs near target genotype-defining SNPs **(Fig. 1)**. EnviroAmpDesigner achieves this by comparing the regions flanking each target genotype-defining SNP against the user-provided whole genome assemblies of non-target organisms **(Supp. Fig. 2)**. These genomes should represent the diversity of taxa expected to be encountered in the samples, especially any closely related organisms as these are most likely to cause off-target amplification (e.g. we recommend including all species of the same family, and all members of related species or serovars/pathovars of the same species, that are not included in the target). We label these as ‘negative genomes’ and the location of the sequences should be specified in the config file as ‘negative_genomes’.

(2)For each set of SNPs selected at the SNP identification stage, EnviroAmpDesigner takes flanking regions upstream and downstream (defined via ‘flank_len_to_check’) of these SNPs from the target reference genome **(Fig. 1)**. For each flanking region, EnviroAmpDesigner searches negative genomes using BLASTn do identify homologues (18). (We tested MMseqs2 as an alternative to BLASTn (19), but due to small size of query file and large number of subject files BLASTn was faster(19).) For each searched reference sequence region, all BLASTn hits are aligned using MAFFT and stored into a Numpy matrix (20, 21). From these matrices, EnviroAmpDesigner identifies positions that are either unique to the target organism or have no homologues in non-target organisms. The organism-specific SNPs identified here, and genotype label specific SNPs selected at the SNP identification stage are the basis of further analysis **(Fig. 1**, **2)**.

### Stage 3: Primer generation

To design primers, EnviroAmpDesigner relies on well-established Primer3, accessed via a python wrapper **(Supp. Fig. 2)** (22, 23). After stage 2 the selected SNPs can fall into three categories. First, genotype-defining SNPs with an organism-defining SNP upstream. Second, genotype label SNPs with an organism SNPs downstream. Third, genotype label SNPs with organism SNPs both upstream and downstream. The first and second cases are mirror versions of each other and are treated the same way, so we will only explain the first of these **(Fig. 1)**. EnviroAmpDesigner treats the third case as two independent instances of first and second, but also generates a pair of primers with each primer’s 3’ end at position of upstream and downstream organism SNPs.

If the organism-defining SNP is upstream of the genotype-defining SNP, the upstream primer is positioned such that the 3’ end is at the organism SNP. This significantly reduces the amplification of non-target organisms thereby improving specificity and sensitivity of target detection (24, 25). To generate downstream primers, the Primer3 software is given the upstream primer and the downstream region (22, 23), the length of which is defined by “*flank_len_to_check*” parameter (suggested default, 1000).

For all downstream primers returned by Primer3, each pair of upstream (fixed) and downstream primer is tested using Primer3 for homo- and hetero-dimerization (22, 23). If a primer pair forms dimers at >10°C, the downstream primer is rejected. (18)We also reject the primers where amplicons are shorter than “*min_amplicon_length”* (suggested default, 200) or longer than double “*flank_len_to_check”* (suggested default, 1000). The remaining primers are output to a tab delimited file with supporting information such as product coordinates, length, and melting temperature.

Finally, EnviroAmpDesigner also uses BLASTn search to identify individual primers with ≥5 nt of 3’ end matching a region in between any pair of primers (18). In our experience, even such short matching sequences are sufficient to cause amplification of truncated amplicons in multiplex PCR **(Fig. 4)**; so we reject such individual primers.

**Figure 4.**
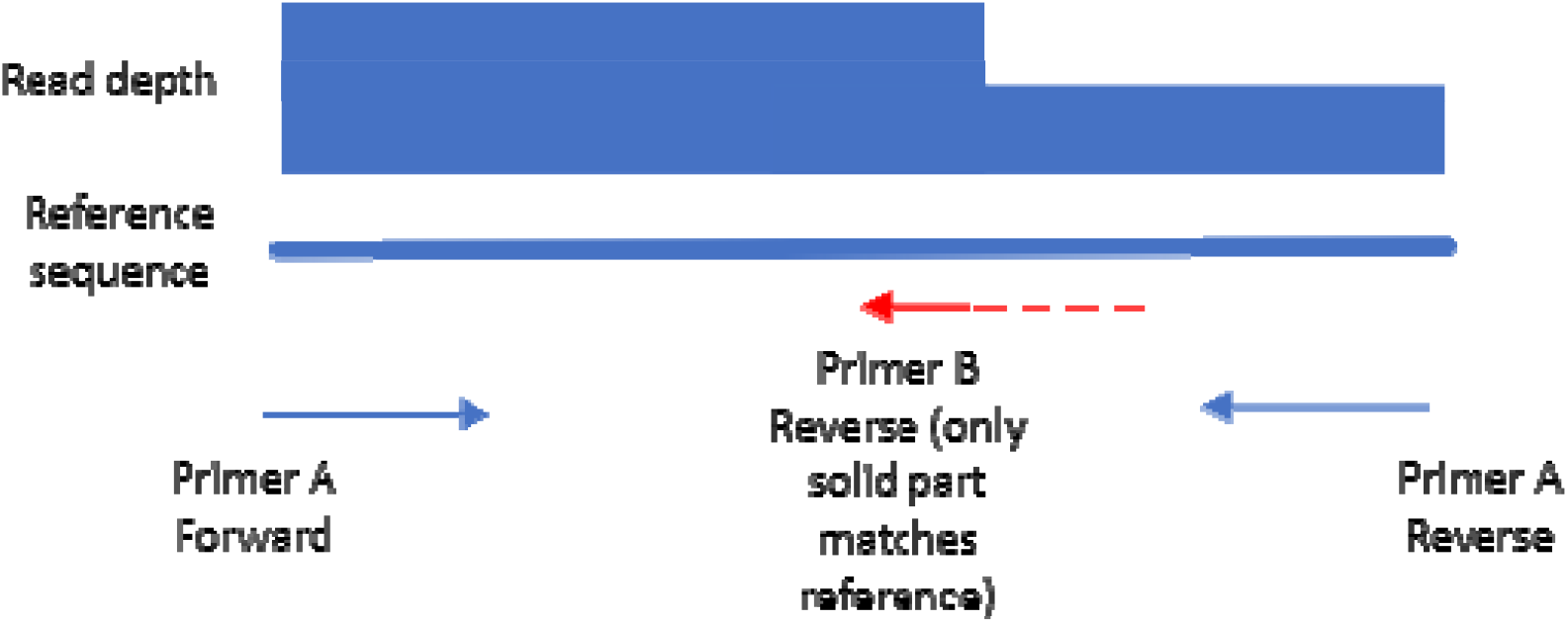
diagram of primer interference in multiplex PCR reaction. Primer B is imperfect match for the reference sequence between primers pair A. However, because it generates shorter amplicon, this compensates for lower annealing efficiency and generates high number of truncated amplicons.

**Figure 5.**
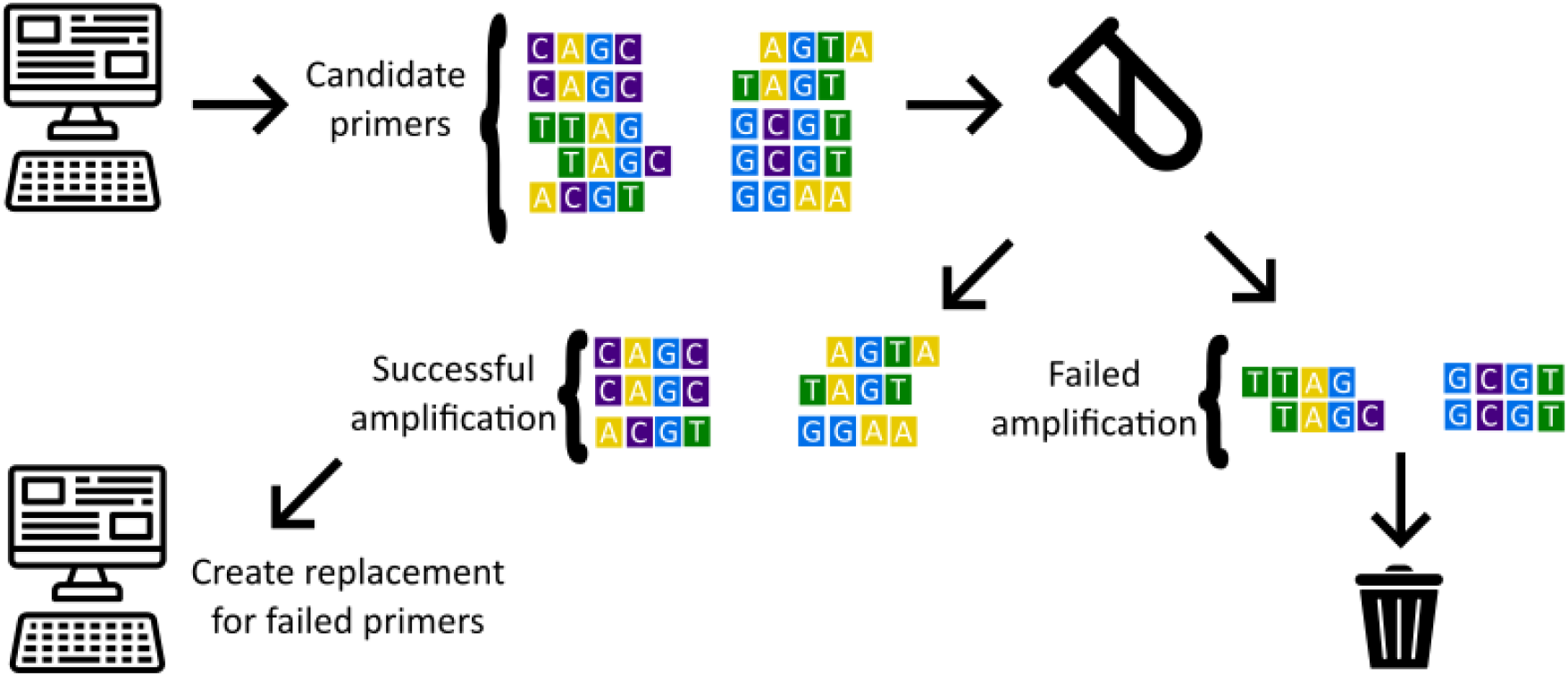
Amplicon primer design workflow with EnviroAmpDesigner. Candidate primers are generated using EnviroAmpDesigner, tested in the lab and for primers that did not produce valid amplification products replacements are generated using EnviroAmpDesigner and further tested in the lab.

**Figure 6.**
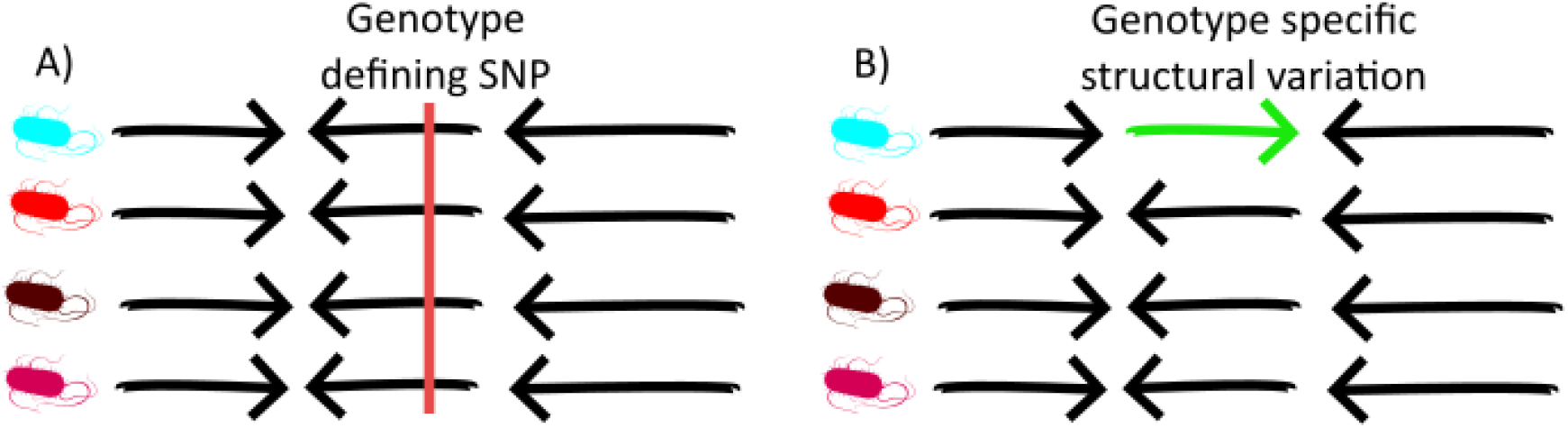
EnviroAmpDesigner will works for SNPs that define target genotype label (A) but is not able to target structural variation (B).

## Results and Discussion

We created EnviroAmpDesigner for design of multiplex amplicon primers for sequencing-based detection and genotyping of pathogens in environmental surveillance samples, but it can also be applied to other samples with large diversity of organisms.

For our motivating use case, we used EnviroAmpDesigner to design a PCR panel for *S*. Typhi targeting 17 genotypes (as defined by the GenoTyphi scheme) (26). For SNP identification (stage 1), we used VCF files for 13,134 *S*. Typhi, generated for a previous population genomics study (1) using a pipeline that mapped Illumina reads against reference genome CT18 (accession AL513382.1) using XYZ (vXYZ) and called variants using bcftools (v1.9)(27). The genotype hierarchy was provided to EnviroAmpDesigner as a **Supp. File 1**. For 4 of the 17 target genotypes, genotype-defining SNPs were found within 1000 bp of SNPs for other genotypes. A total of X genotype-defining SNPs were taken forward for stage 2. To define organism-specific SNPs, we retrieved all reference genomes from the family Enterobacterales (n=202), plus representatives of closely related *S. enterica* serovars *S*. Typhimurium (n=376) and *S*. Paratyphi A (n=20) chosen by randomly downsampling full dataset of NCBI genomes of these organisms (2). For X genotype-defining SNPs (X%), EnviroAmpDesigner identified an organism-specific SNP within 1000 bp. These include X/17 target genotypes (range, X-X candidate regions per genotype), which were taken forward for stage 3 (primer design). For X of these candidate regions (X%), we identified primer pairs that passed all filters. These encompassed X/17 target genotypes. Finally, we selected X primer pairs that provided coverage of genotype-defining SNPs for X/17 genotypes. The config file and genotype hierarchy files available via Zenodo (DOI: 10.5281/zenodo.14967337) **Supp. File 1** and **3**.

We tested the primers using an amplicon sequencing protocol (3) [cite] to generate amplicon data using two S. Typhi isolates (NCBI accessions SRR5949979 and SRR7165748, provided by Satheesh Nair (UKHSA) (4) and a pooled sample consisting of DNA from *S*. Paratyphi A (ATCC 9150D), *S*. Paratyphi B (ATCC-BAA-1250D), *S*. Paratyphi C (ATCC-BAA-1715D), *Aeromonas hydrophila* (ATCC-7965D), *Klebsiella pneumoniae* (ATCC-BAA-1706D), and *Citrobacter freundii* (ATCC-8090D). The resulting amplicon sequences were analysed using our tool AmpliconTyper, as previously reported (28). [X primers failed to XXX, and were redesigned] [summarise data presented in that paper in terms of how consistently the primers yielded amplicons, and callable data for genotype-defining SNPs.]

Based on our experience, we see a typical workflow as an iterative process of primer design using EnviroAmpDesigner, followed by laboratory validation, followed by redesign of the primers for which sequencing produced, in user’s judgement, unsatisfactory number of reads, another round of lab validation and so on until a fully functioning set of primers is achieved.

For the *S*. Typhi panel, we could not identify the cause of underperformance, except for primer-dimer formation which affected X primer sets and motivated us to use stricter thresholds for dimerisation filtering (**Fig. 4**). The iterative nature of primer design motivated us to usilise a single config file to specify all input parameters, as this creates a record of each analysis and simplifies iteration.A typical run with single genotype label target using two target genotypes, 1,293 target organisms VCFs and 331 non-target organisms FASTA genomes takes two minutes using eight CPU cores. As a result, the overall bottleneck in the design process is lab testing. We found that the best way to minimise this bottleneck is to use multiple primer pairs for each target genotype label in second and subsequent design iterations. While this marginally increases the lab work per iteration, it reduces the number of iterations thus shortening the overall design time.

The main limitation of EnviroAmpDesigner is that it cannot design primers for genotypes that are defined by structural variation and do not have any genotype-defining SNPs, as structural variation is not captured by VCFs and thus EnviroAmpDesigner is unaware of it during stage 1 (SNP identification). (22)Structural differences between target and non-target organisms are however captured and used by EnviroAmpDesigner in stage 2. In principle it may be possible to extend EnviroAmpDesigner’s functionality to refine candidate primers targeting genotype-defining structural variants, but this would require extensive development in order to explore edge cases.Another limitation is the potential inaccuracy of reported annealing temperature. While it is possible to explore the full set of Primer3 assumptions to allow user to fine-tune these, we found that lab testing is a more robust approach.

## Conclusions

EnviroAmpDesigner offers a robust tool for design of multiplex PCR primers for detection and typing of target organisms from environmental or similarly complex samples. It relies on standard input files, includes a range of filtering steps to improve reliability of generated primers and can run be run on a mid-range desktop computer. The tool is freely available via Conda and GitHub (29).

## Abbreviations

ATCC: American type culture collection
AMR: antimicrobial resistance
BAM: Binary Alignment Map
NCBI: National Center for Biotechnology Information
ONT: Oxford Nanopore Technologies
PCR: Polymerase chain reaction
RT-qPCR: quantitative reverse transcription polymerase chain reaction
SNP: single nucleotide polymorphism
UKHSA: United Kingdom Health Security Agency
VCF: Variant Call File
WGS: whole genome sequencing

## Availability and requirements

Project name: EnviroAmpDesigner

Project home page: https://github.com/AntonS-bio/EnviroAmpDesigner.git

Installation: via Conda package EnviroAmpDesigner

Operating system(s): Linux, MacOS or Windows 10+ with Windows Subsystem Linux

Programming language: Python

Other requirements: None

License: GNU GPL v3

Any restrictions to use by non-academics: None

## Ethics approval and consent to participate

Not applicable.

## Consent for publication

Not applicable.

## Authors’ contributions

Conceptualization: NG, KEH. Data curation: DA, BB, KG, VRM, ZD. Formal analysis: AS, ZD, JM, CT. Funding acquisition: NG, KEH, DA. Investigation: JM, CT, MO, YAS, EOD, DA, BB, KG, VRM. Software: AS. Resources: All authors. Writing – original draft: AS. Writing – review & editing: All authors.

---

**Supplementary Figure 1.**
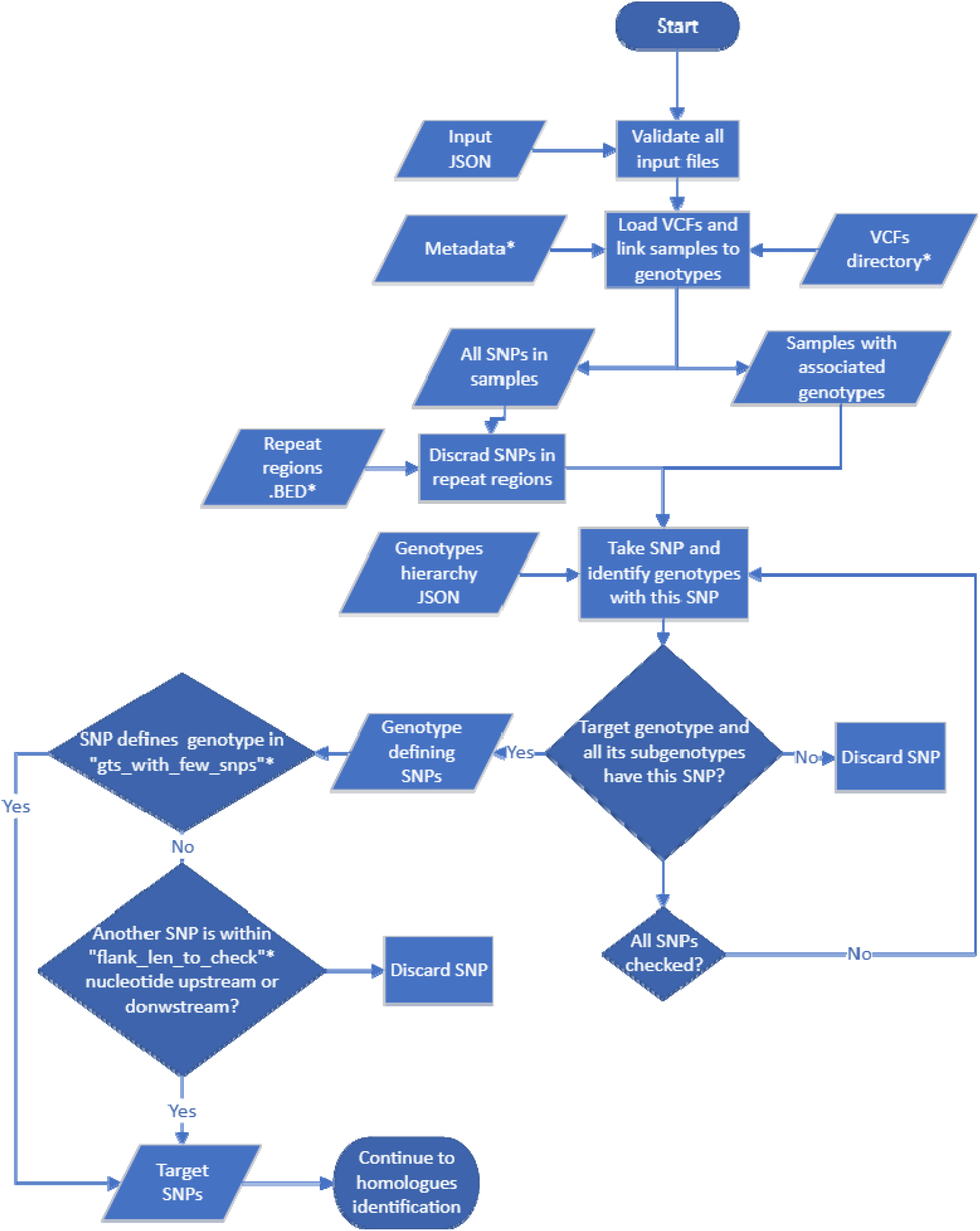
identification of genotype-defining SNPs.

**Supplementary Figure 2.**
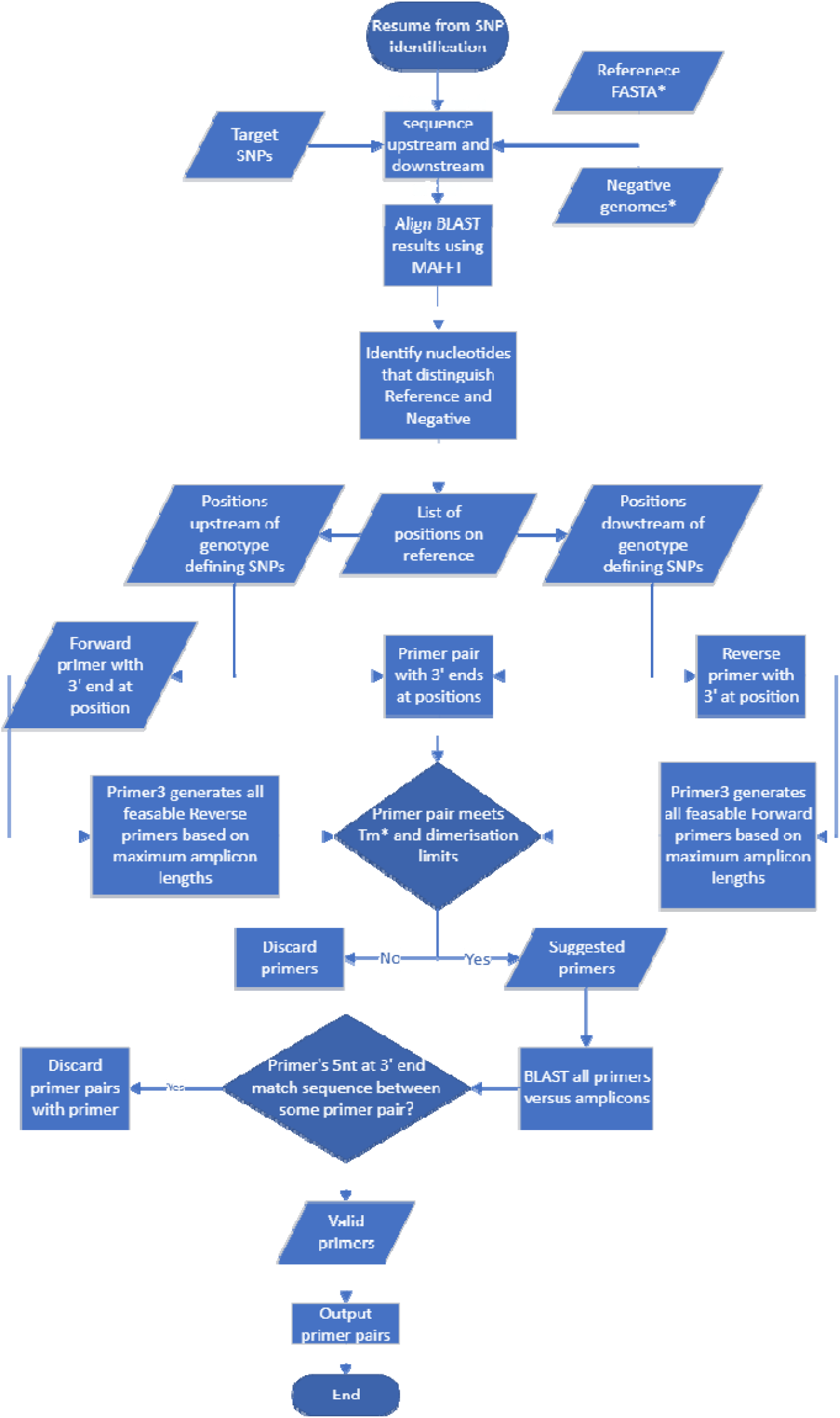
identification of target organism defining SNPs and primer design.

## References

1. Carey ME, Dyson ZA, Ingle DJ, Amir A, Aworh MK, Chattaway MA, et al. Global diversity and antimicrobial resistance of typhoid fever pathogens: Insights from a meta-analysis of 13,000 Salmonella Typhi genomes. Elife. 2023;12.

2. Agarwala R, Barrett T, Beck J, Benson DA, Bollin C, Bolton E, et al. Database resources of the National Center for Biotechnology Information. Nucleic Acids Research. 2018.

3. Mahindroo J, Spadar A, Troman C, Dyson Z, Holt K, Grassly N. Amplicon Sequencing for Genotyping S. Typhi V.3 DOI: 10.17504/protocolsio36wgq31dylk5/v3. 2024.

4. Ashton PM, Nair S, Peters TM, Bale JA, Powell DG, Painset A, et al. Identification of Salmonella for public health surveillance using whole genome sequencing. PeerJ. 2016;4:e1752.

5. Grassly NC, Shaw AG, Owusu M. Global wastewater surveillance for pathogens with pandemic potential: opportunities and challenges. Lancet Microbe. 2025;6(1):100939.

6. Klapsa D, Wilton T, Zealand A, Bujaki E, Saxentoff E, Troman C, et al. Sustained detection of type 2 poliovirus in London sewage between February and July, 2022, by enhanced environmental surveillance. Lancet. 2022;400(10362):1531–8.

7. Navarro A, Gomez L, Sanseverino I, Niegowska M, Roka E, Pedraccini R, et al. SARS-CoV-2 detection in wastewater using multiplex quantitative PCR. Sci Total Environ. 2021;797:148890.

8. Abraham D, Kathiresan L, Sasikumar M, Aiemjoy K, Charles RC, Kumar D, et al. Wastewater surveillance for Salmonella Typhi and its association with seroincidence of enteric fever in Vellore, India. PLoS Negl Trop Dis. 2025;19(3):e0012373.

9. Owusu M, Darko E, Akortia D, Nkrumah G, Twumasi-Ankrah S, Owusu-Ansah M, et al. Evaluation of Moore and grab sampling method for Salmonella Typhi detection in environmental samples in Ghana. PLoS One. 2025;20(2):e0318840.

10. Srivathsan A, Meier R. Scalable, Cost-Effective, and Decentralized DNA Barcoding with Oxford Nanopore Sequencing. Methods Mol Biol. 2024;2744:223–38.

11. de Cesare M, Mwenda M, Jeffreys AE, Chirwa J, Drakeley C, Schneider K, et al. Flexible and cost-effective genomic surveillance of P. falciparum malaria with targeted nanopore sequencing. Nat Commun. 2024;15(1):1413.

12. Farooqui A, Khan A, Kazmi SU. Investigation of a community outbreak of typhoid fever associated with drinking water. BMC Public Health. 2009;9:476.

13. Qamar FN, Yousafzai MT, Khalid M, Kazi AM, Lohana H, Karim S, et al. Outbreak investigation of ceftriaxone-resistant Salmonella enterica serotype Typhi and its risk factors among the general population in Hyderabad, Pakistan: a matched case-control study. Lancet Infect Dis. 2018;18(12):1368–76.

14. Matrajt G, Lillis L, Meschke JS. Review of Methods Suitable for Environmental Surveillance of Salmonella Typhi and Paratyphi. Clin Infect Dis. 2020;71(Suppl 2):S79–S83.

15. Hendling M, Pabinger S, Peters K, Wolff N, Conzemius R, Barisic I. Oli2go: an automated multiplex oligonucleotide design tool. Nucleic Acids Res. 2018;46(W1):W252–W6.

16. Gervais AL, Marques M, Gaudreau L. PCRTiler: automated design of tiled and specific PCR primer pairs. Nucleic Acids Res. 2010;38(Web Server issue):W308–12.

17. Thomsen MCF, Hasman H, Westh H, Kaya H, Lund O. RUCS: rapid identification of PCR primers for unique core sequences. Bioinformatics. 2017;33(24):3917–21.

18. Altschul SF, Madden TL, Schäffer AA, Zhang J, Zhang Z, Miller W, et al. Gapped BLAST and PSI-BLAST: A new generation of protein database search programs. Nucleic Acids Research1997. p. 3389–402.

19. Steinegger M, Soding J. MMseqs2 enables sensitive protein sequence searching for the analysis of massive data sets. Nat Biotechnol. 2017;35(11):1026–8.

20. Katoh K. MAFFT: a novel method for rapid multiple sequence alignment based on fast Fourier transform. Nucleic Acids Research. 2002;30(14):3059–66.

21. Harris CR, Millman KJ, van der Walt SJ, Gommers R, Virtanen P, Cournapeau D, et al. Array programming with NumPy. Nature. 2020;585(7825):357–62.

22. Untergasser A, Cutcutache I, Koressaar T, Ye J, Faircloth BC, Remm M, et al. Primer3--new capabilities and interfaces. Nucleic Acids Res. 2012;40(15):e115.

23. Pruitt BC N., Primer3-py. 0.22.0 ed 2014.

24. Simsek MA H., Effect of single mismatches at 3′–end of primers on polymerase chain reaction. Journal of Scientific Resaerch - Medical Sciences. 2000;2(1):11–4.

25. Apte A, Daniel S. PCR primer design. Cold Spring Harb Protoc. 2009;2009(3):pdb ip65.

26. Dyson ZA, Holt KE. Five Years of GenoTyphi: Updates to the Global Salmonella Typhi Genotyping Framework. J Infect Dis. 2021;224(12 Suppl 2):S775–S80.

27. Danecek P, Bonfield JK, Liddle J, Marshall J, Ohan V, Pollard MO, et al. Twelve years of SAMtools and BCFtools. GigaScience. 2021;10(2).

28. Spadar A, Mahindroo J, Troman C, Owusu M, Adu-Sarkodie Y, Owusu-Dabo E, et al. AmpliconTyper – tool for analysing ONT multiplex PCR data from environmental and other mixed sources. bioRxiv. 2025.

29. Anaconda. v3.0.6 ed: Andaconda Inc; 2024.

